# Design and demonstration in vitro of a mouse-specific Transcranial Magnetic Stimulation coil

**DOI:** 10.1101/2020.01.09.900993

**Authors:** Farah A. Khokhar, Logan J. Voss, D. Alistair Steyn-Ross, Marcus T. Wilson

## Abstract

**Background:** Transcranial Magnetic Stimulation (TMS) is a technique used to treat different neurological disorders non-invasively. A pulsed current to a coil generates a magnetic field (B-field) which induces an electric field (E-field). Underlying biophysical effects of TMS are unclear. Therefore, animal experiments are needed; however, making small TMS coils suitable for mice is difficult and their field strengths are typically much lower than for human sized coils.

**Objectives/Hypothesis:** We aimed to design and demonstrate a mouse-specific coil that can generate high and focused E-field.

**Methods:** We designed a tapered TMS coil of 50 turns of 0.2 mm diameter copper wire around a 5 mm diameter tapered powdered iron core and discharged a 220 μF capacitor at 50 V through it. We measured B-field with a Hall probe and induced E-field with a wire loop. We measured temperature rise with a thermocouple. We applied 1200 pulses of continuous theta burst stimulation (cTBS) and intermittent theta burst stimulation (iTBS) to mouse brain slices and analysed how spontaneous electrical activity changed.

**Results:** The coil gave maximum B-field of 685 mT at the base of the coil and 340 mT at 2 mm below the coil, and maximum E-field 2 mm below the coil of approximately 10 V/m, at 50 V power supply, with a temperature increase of 20 degrees after 1200 pulses of cTBS. We observed no changes in B-field with heating. cTBS reduced frequency of spontaneous population events in mouse brain slices up to 20 minutes after stimulation and iTBS increased frequency up to 20 minutes after stimulation. No frequency changes occurred after 20 minutes. No changes in amplitude of spontaneous events were found.

**Conclusion:** The design generated fields strong enough to modulate brain activity in vitro.

## I. Introduction

Transcranial Magnetic Stimulation (TMS) is a non-invasive medical technique [1, 2] used in treatment of major depression [3–5] and therapeutically tested for epilepsy [6], stroke [2], Parkinson’s disease [7], functional hand dystonia [8–10] and Alzheimer’s disease [11–14]. During TMS, a large pulse of current of less than 1 millisecond duration passes through a TMS coil placed on the head. This current pulse produces a changing magnetic field which induces an electric field within the brain [15–19]. While the physics of the coil is well-understood, the fundamental neurophysiological effects of the induced electric field, such as TMS-induced plasticity [20–22], changes in cortical excitability [20, 23–26] and TMS-induced neuronal effects [27], are not clear. There is a need to perform invasive measurements to obtain a profound understanding of the underlying principles of TMS [24–33]. Therefore, animal experiments are needed to obtain a better understanding behind the mechanisms of TMS-induced plasticity and changes in excitability. Mice are particularly suitable for invasive experiments because their brain geometry and connections have been well mapped [34].

Classic figure-of-eight human TMS coils, around 60 – 70 mm diameter [35, 36], with magnetic field strength of 1.5 – 2.5 T [15, 36–38] and induced electric field strength of 150 – 250 V/m [39] are designed to stimulate a small and focused area of human cortex. Applied to a mouse, the human TMS coil would stimulate the entire mouse brain. Therefore, a miniaturized coil is required that could mimic the electromagnetic field strengths of a human TMS coil [18, 19, 28, 35]. The mouse-specific TMS coil should be designed in terms of the size difference of human and mouse brain. Broadly, the mouse brain is smaller than the human brain by a factor of approximately 10 in linear dimensions [40]. Additionally, the surface of the mouse brain lies approximately 2 mm below the skin surface at the top of the head [41] compared to human brain which is around 20 mm below from the surface [42]. Therefore, one would expect an appropriate mouse coil to be of order 6 – 7 mm in size. There are several physical and technical challenges in scaling down to small-scale TMS coils. For example, applying large currents will lead to overheating [19, 28, 43, 44]. Furthermore, there is a lack of clarity on what parameters should be optimised in a small-scale coil design [19, 45]. Options could include greater stimulation focality by offsetting coil position [46, 47], improving stimulation focality by scaling down human TMS coils [48], or achieving high magnetic field intensities [49]. In general, there is a trade-off between greater electric field intensity and focality [50], which makes design of mouse coils particularly challenging.

Optimizing both magnetic field and electric field simultaneously may be unachievable. Since curl ***E*** = −*∂**B***/*∂t*, the dimensions of electric field ***E*** and the time-derivative of magnetic field ***B*** differ by a spatial term and so magnetic and electric field strengths do not scale with size in the same way [19]. To scale the magnetic field intensity requires only one-tenth current in a tenth-size coil to match the intensity of a human TMS coil, whereas to match the electric field intensity requires the same rate of change of current as for human coils. Assuming that the time scale for the pulses are the same for the mouse and human coils, this requires similar current levels for the mouse and human coils, which is likely to create hazardous levels of Joule heating in a mouse coil [1, 51].

There has been some previous work on small coil design. Tang et al. constructed an iron-core rodent-specific TMS coil having 8 mm outer diameter by winding 780 turns of insulated copper wire that delivered the magnetic field strength of approximately 120 mT and an induced electric field strength modelled at around 12.7 V/m on the surface of a rat cortex at 100 V power supply [28]. Wilson et al. have designed and constructed a 5 mm mouse-specific ferrite core coil. The ferrite eliminated eddy currents that were present in the Tang et al. core [28]. Windings were reduced to 70 turns to reduce inductance. The coil demonstrated a magnetic field strength of around 180 mT, at a depth below 2 mm below the coil, the induced electric field was estimated at around 2.5 V/m in air for a 30 V power supply [19]. A similar-sized coil for peripheral-nerve stimulation has also been demonstrated in a figure-of-eight configuration by Colella et al. [52]. However, much smaller coils have also been produced. For example, Bonmassar et al. have demonstrated that a sub-millimeter coil (diameter 0.5 mm), with B-fields of magnitude 10 mT and Efields around 6 V/m, can change action potential production in retinal ganglion cells [53]; these coils are also safe for use with magnetic resonance imaging (MRI), raising temperature by just 1 °C under a MRI protocol at 1.5 T and 64 MHz [54]. Minusa et al. used an implantable set of magnetic stimulators to apply magnetic stimulation directly to the brain surface of mice. Coils were 1 mm long and 0.5 mm diameter and provided a highly-focused *B*-field of around 5 mT [55]. Despite this recent progress, these fields remain much lower than the fields of equivalent human TMS coils [56]. Coils do not need to be cylindrically symmetric. Indeed, departure from this symmetry restriction opens up novel possibilities. For example, Meng et al. demonstrated that tilting the axis of a coil’s core, with respect to the coil’s axis can result in a focused region of increased field [57]. Sanchez et al. have used a stream function method to design coils for focused unilateral hemispheric stimulation of a rodent brain while minimizing heating [58].

In this work, we have designed a mouse-specific coil with increased field strengths. To demonstrate its effect, we have applied 1200 pulses of continuous theta-burst stimulation (cTBS) and intermittent theta-burst stimulation (iTBS) [59] to mouse brain slices in vitro. Theta-burst stimulation consists of application of short bursts of pulses (usually 3 pulses in a burst, 20 ms apart, with 5 bursts per second) and is often used to investigate plasticity and excitability mechanisms within the brain. In cTBS experiments, we have applied bursts continuously for 80 seconds, whereas in iTBS experiments we have repeated 2 seconds of stimulation followed by 8 second pauses for 400 seconds.

We have used spontaneous electrical activity in the cortex, measured through the local field potential, as a simple and readily demonstrable measure of the impact of stimulation. Specifically, we measure the amplitude and rate of ‘seizure-like events’ (SLEs), or spontaneous epileptiform-like activity as shown in Figure 1, due to correlated firing of groups of neurons in a slice that is prepared in an artificial cerebrospinal fluid from which magnesium ions have been removed [60]. This is an established and straightforward method for quantifying N-Methyl-D-aspartic acid (NMDA)-dependent excitability (including long-term potentiation) of brain tissue [61, 62]. While the method does not provide subtle detail about any mechanisms of TMS interaction with the brain, it does serve to allow a simple demonstration that the field strengths are sufficient for unspecified changes in the brain to occur as a result of stimulation.

**Fig. 1.**
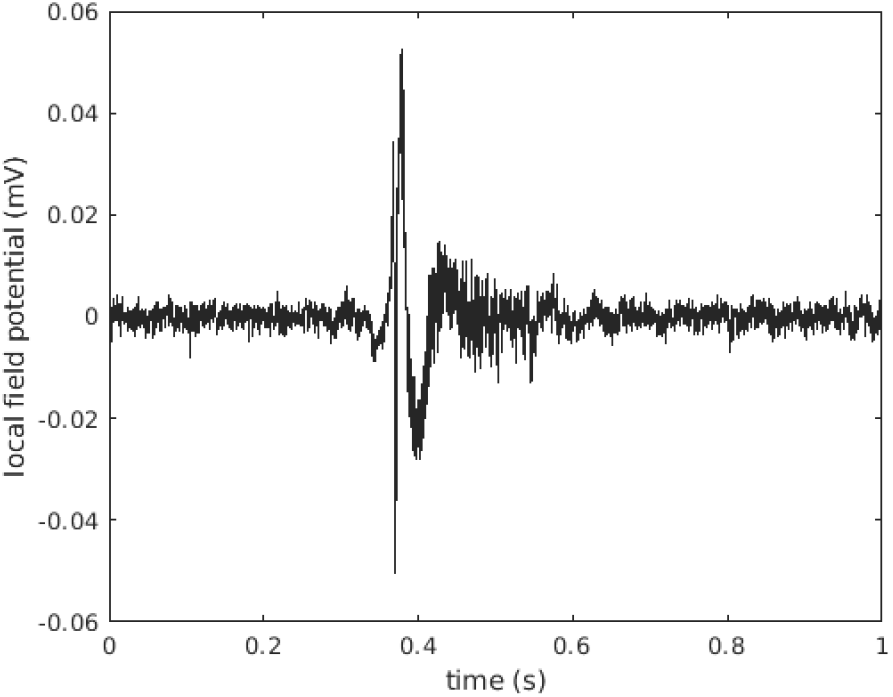
Typical example of an expanded view of a single no-magnesium seizure-like event.

We begin by describing the coil design. We then give details of the methodology for measuring the coil performance in terms of field strengths and spatial distribution, and the effect on SLEs in vitro. Finally, we present the results and discuss implications for use of TMS coils for mice and suggest future coil designs.

## II. Methods

### A. Coil Design

We have constructed a mouse-specific coil by winding 50 turns of copper wire of conductive diameter 0.4 mm onto a 5 mm diameter carbonyl powdered iron core (Micrometals, U. S. A). The windings are secured in place with glue (Loctite^®^ Superglue). This was adequate to ensure that coil windings did not come loose. A larger diameter wire than used in [19] allows the coil to have a much lower resistance, increasing the current. To reduce inductance, we have reduced the windings from 70 (as used in [19]) to 50. The smaller number of turns decreases the inductance in the coil and allows a rapid change in current, which will increase the *E*-field strength [19]. We used carbonyl powdered iron core (Micrometals, U.S.A) as a substitute to the soft-ferrite core [19] since it has a higher saturation magnetization (about 1.5 T) than ferrite (about 0.4 T). However, powdered iron cores have lower permeability than ferrite due to their air-gap distribution. The length of the untapered powdered iron core was *A* = 19 mm and the diameter *B* = 5 mm. In an attempt to increase focality of the fields, we tapered the core over the final *C* = 3 mm of its length to a diameter of *D* = 2 mm, by careful sanding. We also followed a tapered design for the coil to focus fields on a small area while giving a high field strength. The coil’s height was *E* = 12 mm and its diameter at the non-tapered end was *F* = 9 mm. It was tapered over its final *G* = 3 mm of length to a diameter of just *H* = 4 mm. The coil is shown in Figure 2. The electrical parameters of the coil such as resistance and resistance of the coil against frequency was measured with an Agilent E4980A four-point impedance meter (Agilent Instruments, Santa Clara, California, USA). During the design and prototyping process we made several coils, although only one was tested in detail and reported below.

**Fig. 2.**
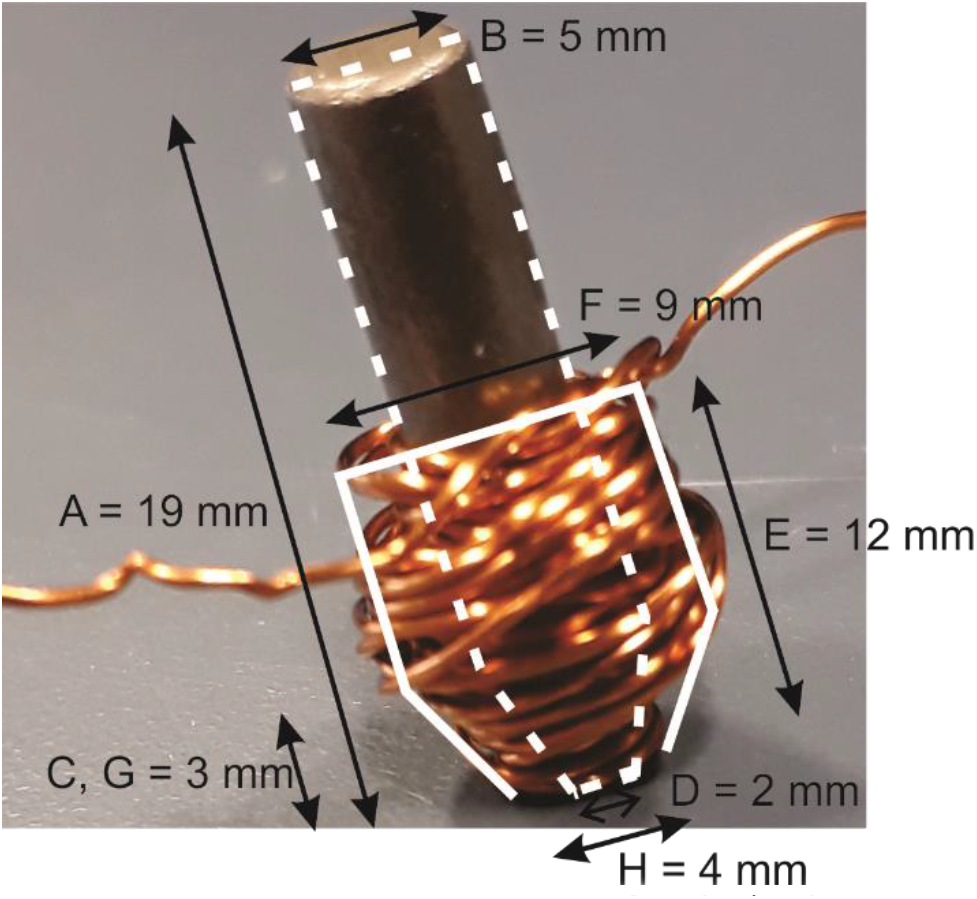
A photograph of the coil. 50 turns of conductive diameter of copper wire (0.4 mm). The dimensions of the core and coil are shown on the figure. The solid line shows the approximate shape of the coil and the dashed line the shape of the core.

The cylindrical geometry implies that the B-field has no azimuthal component, and that the field is rotationally symmetric. Mathematically, ***B*** = *B_z_*(*z, r*) ***a**_z_* + *B_r_*(*z, r*) ***a***_*r*_, where *z* is the on-axis distance below the base of the coil, *r* is the radial distance from the axis and ***a**_z_* and ***a**_r_* are unit vectors in the axial and radial directions respectively. On-axis, where *r* = 0, the field is purely axial, that is ***B*** = *B_z_*(*z*) **a**_*z*_. With cylindrical symmetry the induced electric field is purely azimuthal, that is ***E*** = *E*_θ_(*z, r*) ***a***_θ_ where ***a***_θ_ is a unit vector in the azimuthal direction.

We created a TMS pulse by discharging an electrolytic capacitor (nominal value 220 μF ± 20%, rated to 50 V, Panasonic) at 50 V through the coil. The capacitance was measured with the Agilent E4980A four-point impedance meter across the range 20 Hz to 2 MHz. At frequencies below 10 kHz capacitance was constant at 206 μF. The self-resonant frequency was found to be approximately 60 kHz, well above the frequency scales for coil discharge. The circuit was closed by gating an AUIRL3705N low on-resistance (10 mΩ) Metal-Oxide Semiconductor Field Effect Transistor (MOSFET) at 10 V, allowing rapid discharge of the capacitor through the coil. The gate voltage was provided by boosting the output from an Arduino Uno microcontroller with an operational amplifier. For each pulse, the gate was held open for 2 ms, which was long enough for oscillations in the LC circuit to die away. The Arduino can be programmed to provide specific stimulation protocols. A circuit diagram is shown in Fig. 3. While more complicated implementations exist, for example for controlling pulse shape, the circuit is sufficient for applying a high current pulse to the coil. In this work we have used two protocols: 1200 pulses (80 seconds) of continuous theta burst stimulation at 3 pulses per burst and 5 bursts per second, and 1200 pulses (400 seconds) of intermittent theta burst stimulation at 3 pulses per burst and 5 bursts per second repeated for 2 seconds ON and 8 seconds OFF.

**Fig. 3.**
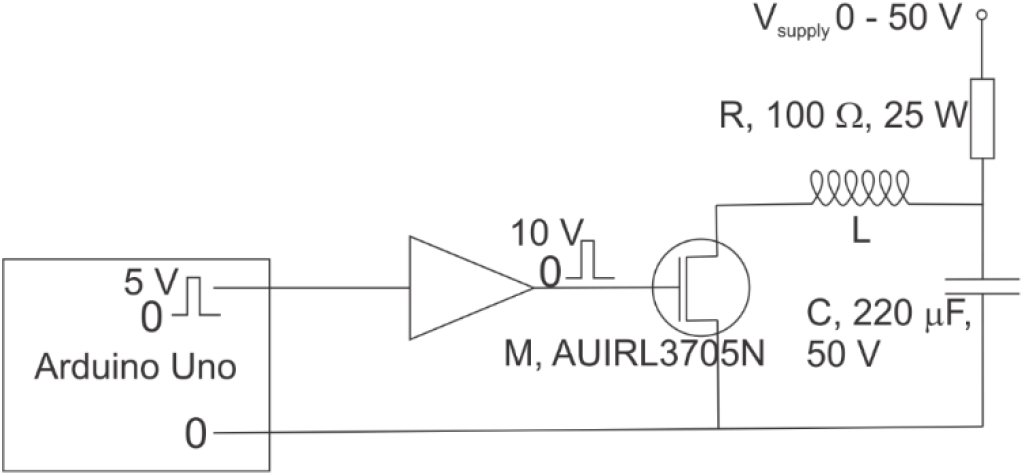
The circuit diagram. When the MOSFET M is open, the capacitor C charges via the resistor R up to the supply voltage. When M closes, the capacitor discharges through the coil L. The gating of M is provided with 10 V pulses in cTBS or iTBS sequences, provided by the amplified output of an Arduino Uno microcontroller.

**Fig. 4.**
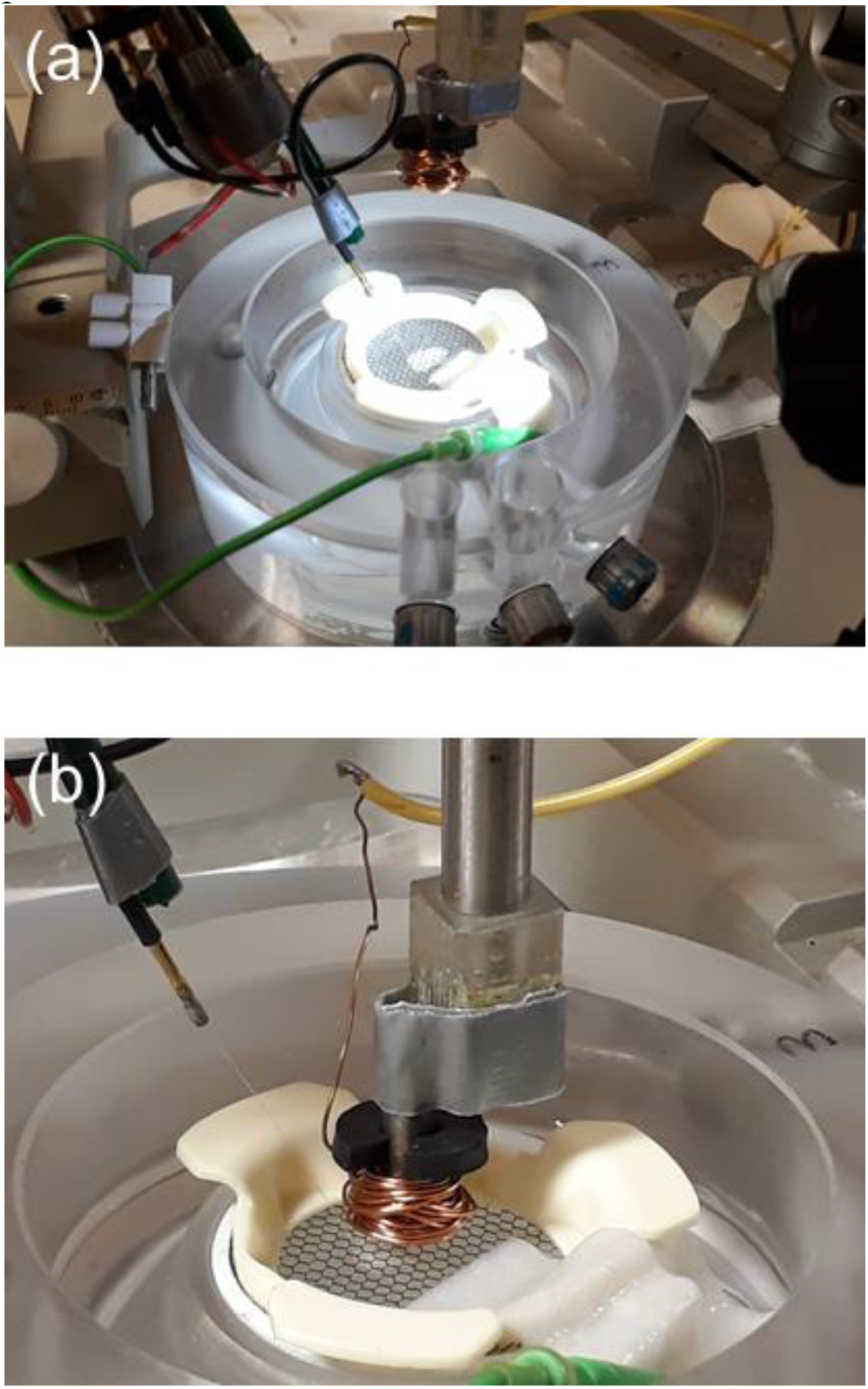
The set-up for the in vitro experiments. (a) The slice with recording electrode in place. (b) The coil in position for TMS application 2 mm above the slice.

### B. Coil Modeling

We have modelled the electromagnetic fields of the coil using COMSOL Multiphysics with 2-D axial symmetry. The maximum current in the 50 turns of the coil was estimated to be at most 140 A by assuming that the electrostatic energy in the 220 microfarad capacitor at 50 V is completely transferred to magnetic energy in the coil, using the measured value of the coil inductance (27.1 microhenry). A frequency-domain calculation with frequency equal to 1800 Hz was used, corresponding to the experimentally-measured biphasic pulse period of about 0.55 ms. The geometry of the modelled coil was slightly simplified by modelling the boundaries between the core and air, windings and air, and core and windings as straight line segments. However, the turn density in the COMSOL simulation was identical to the turn density of the actual coil. The core was modelled as iron, but with the electrical conductivity set to zero to account for powdering of the core eliminating eddy currents. The coil was modelled as copper, but with the axial and radial electrical conductivities set to zero to prevent modelling of currents in these directions.

### C. Measurement of B- and E-fields

The axial component of the *B*-field, *B_z_*, was measured as a function of axial distance *z* and radial distance from the axis, *r*, using an A1302KUA-T Hall effect sensor (ALLEGRO MICROSYSTEMS). On the axis of the coil, the B-field is expected to be purely axial and the Hall probe was oriented accordingly. The sensor was chosen since it gives high spatial and sufficient temporal resolution (output bandwidth 20 kHz) for B-field, for fields up to order 200 mT in magntitude. The Hall sensor was calibrated using a Helmholtz coil arrangement; known currents were applied to the coil, the B-field at the centre of the Helmholtz coil calculated from the current and geometry, and the Hall voltage measured. The sensitivity of the sensor, in terms of B-field per unit voltage, was found to be 90(6) mT/V, in agreement with the manufacturer’s specification of 77 mT/V ± 20%. The sensor measures a maximum field of 225 mT. While this is sufficient for much of the region of space in the vicinity of the coil, very close to the coil the fields were larger at the highest supply voltages we used. To infer the B-field at the base of the coil we used measurements made further from the coil, where fields are weaker, as described in the appendix.

The corresponding *E*-field was measured as a function of radial distance, 2 mm below the coil, by measuring the voltage induced around a thin loop of wire. This is equivalent to measuring the rate of change of flux through the loop. The cylindrical symmetry of the coil allows the measurements to be carried out with a single orientation; a full three-dimensional mapping of electric field direction is not required since the E-field is purely azimuthal. A thin loop of wire was oriented in such a way that the long sides were perpendicular to the *E*-field direction, and the short side (length 1.5 mm) parallel to the direction [39]. The geometry is shown in more detail in the appendix. The induced voltage around the loop divided by the length of the short side gives the approximate *E*-field. Such an arrangement measures only induced electric field, not static electric field, but we would expect with a cylindrically symmetric coil in air the latter to be zero. Our measurements are not, therefore, a direct indication of the *E*-field to be expected in the vicinity of the mouse brain, where charge will build-up at interfaces between regions of different conductivity. However, they are still of use in terms of comparing between different coil designs. We have chosen 2 mm distance below the coil because it approximates the distance from the coil to the mouse cortex in mice experiments [26].

### D. Measurement of Temperature

A thermocouple temperature probe was inserted against core before the coil was wound. This allowed temperature of the core and coil assembly to be recorded. The temperature was monitored during the application of 1200-pulse cTBS and iTBS protocols, with the coil in air.

### E. In Vitro Experiments

The brain slice experiments used for this research were approved by the Animal Ethics Committee at the University of Waikato (Hamilton, New Zealand). The experiments were performed in a Faraday shielded room to control electrical noise interference. Details of the experimental method for preparing slices in artificial cerebrospinal fluid (aCSF) and measuring electrophysiological activity are given in the appendix.

We analysed the spontaneous seizure-like events (SLEs) in terms of amplitude and inter-event frequency. The amplitude of an SLE is defined as the maximum (peak) value of the LFP minus the minimum (trough) value during the course of a single SLE. The inter-event frequency is defined as the number of events in a given time interval (in the case of this work, 10 minutes) divided by the time interval. Of the 20 slices, 10 were used for cTBS and 10 for iTBS experiments. Each slice received both stimulation and sham treatments, but half received stimulation first, and half received sham first. For stimulation, the coil was clamped 2 mm above the slice (touching the slice perfusion fluid). For sham treatment the coil was clamped 20 mm above the slice and thus received much lower EM field strengths. For the stimulation-first slices, the timeline consisted of 15 minutes baseline recording of SLEs, then 1200 pulses of TBS (either cTBS or iTBS), then 30 minutes of SLE recording. This was followed by 1200 pulses of sham stimulation (cTBS or iTBS) and 30 more minutes of SLE recording. Figure 5 parts (a) and (c) show these sequences for cTBS and iTBS respectively. For the sham-first slices, the timeline was the same, except that the order of sham and stimulation was reversed. These sequences are shown in Figure 5 parts (b) and (d) for cTBS and iTBS respectively. The application of the TMS to the slice resulted in considerable interference to the electrophysiological voltage recording. However, we emphasize that the recording became stable approximately a second after the end of the TMS period and any artefact, where it existed, was excluded from the analysis. The electrophysiological recording equipment provides almost no current to the slice, and thus is unlikely to change the time course of the applied TMS pulses.

**Fig. 5.**
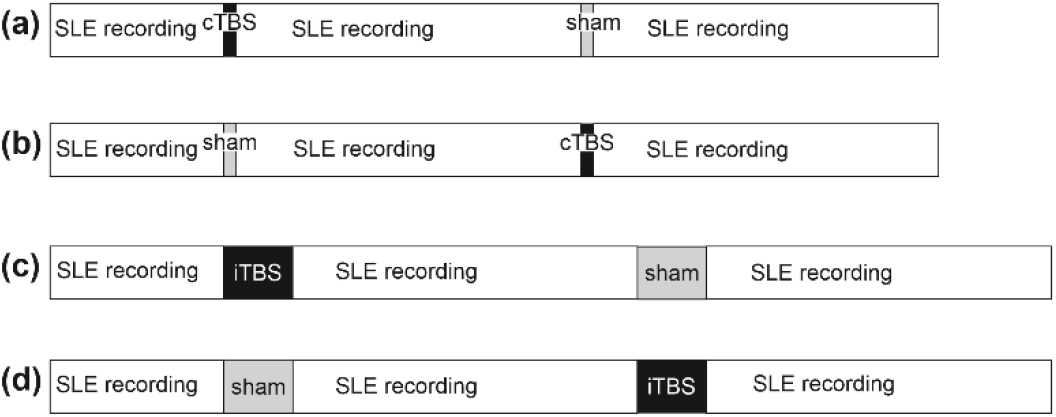
The timeline of the protocols. (a) For the cTBS experiments, activity from five slices was recorded for 15 minutes (baseline), then the slice was subjected to 1200 pulses of cTBS. The activity was then recorded for 30 minutes, before a sham stimulation and 30 minutes recording. (b) Also for cTBS, five more slices were stimulated as for (a), but with the order of cTBS and sham stimulation reversed. (c) and (d) For the iTBS experiments, the timeline was as for cTBS, except that the cTBS stimulation was replaced with iTBS. Five slices were stimulated with iTBS first, and another five with sham first.

Post-stimulation and post-sham LFPs were analysed in three 10-minute segments, specifically 0-10 min, 10-20 min and 20-30 min after stimulation or sham. Average SLE amplitude and inter-event frequency were computed in each segment and the relative changes compared with the baseline were calculated. For the second stimulation in a protocol (i.e. the sham stimulation of Figure 5 (a) and (c), or the cTBS and iTBS of Figure 5 (b) and (d) respectively) the activity in the 10-minute segment immediately before the second stimulation was used as the baseline.

## III. RESULTS

### A. Electrical properties of the coil

Figure 6(a) shows the resistance of the coil as a function of frequency. It is constant at about 212 mΩ for frequencies less than around 10 kHz and begins to rise for higher frequencies. We are particularly interested in frequencies of around 10 kHz since the timescale of the TMS pulse is the order of 0.1 ms and therefore the increased resistance due to higher frequencies is unlikely to be a problem [19]. The reactance of the coil shows a linear behaviour with frequency, shown in Figure 6(b). The inductance of the coil is constant at about 27.1 μH across the frequency range.

**Fig. 6.**
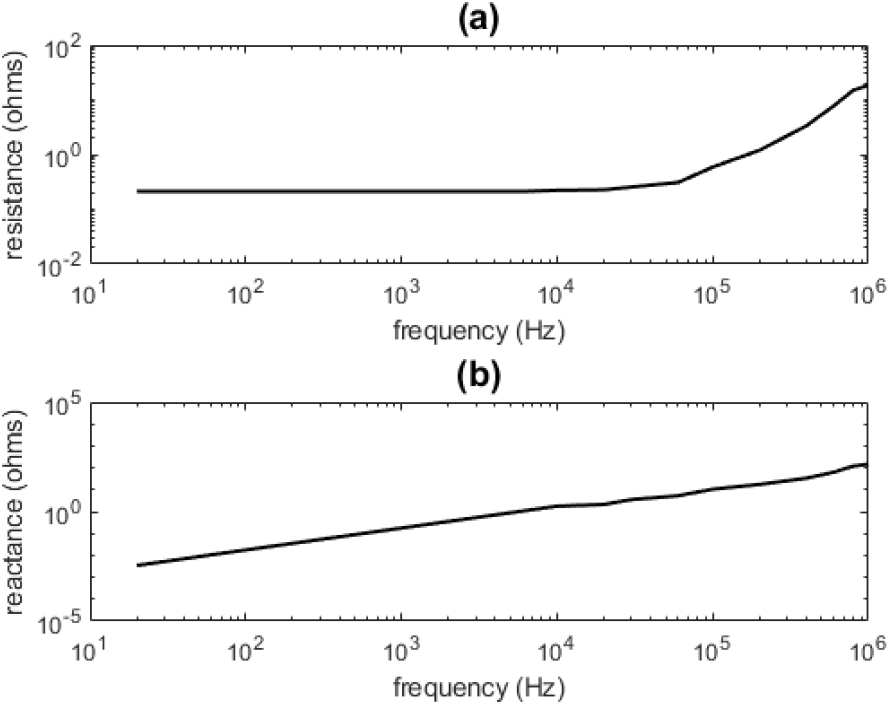
The electrical properties of 50 turn tapered powdered iron coil as measured with an Agilent E4980A four-point impedance meter. (a) The resistance of the 50 turn tapered powdered iron coil as a function of frequency. (b) The reactance of the 50 turn tapered powdered iron coil as a function of frequency.

### B. B-field and E-field

A measurement of the *B*-field strength as a function of supply voltage up to 50 V is shown in Fig. 7. The black line shows measurements at the base of the coil (*z*=0); the solid blue line shows on-axis measurements at *z*=3 mm below the coil. The dashed blue line shows the *z*=3 mm measurements scaled to match the *z*=0 mm measurements at the lower voltages; the two curves match very well and the continuation of the dashed blue curve then indicates an estimate of B-field at the base of the core at the higher voltages. Seven such measurements were taken, with the mean B-field at 50 V supply being measured as 685 mT with standard error in the mean of 35 mT. The uncertainty is attributed to small errors in positioning the sensor with respect to the coil. The B-field grows approximately linearly with supply voltage for voltages up to around 20 V, before reducing in its response. However, at 50 V supply, the field still shows significant increase in strength with increasing voltage, suggesting that the core has still not fully saturated. The B-field showed no change when the temperature was raised by 20°C.

**Fig. 7.**
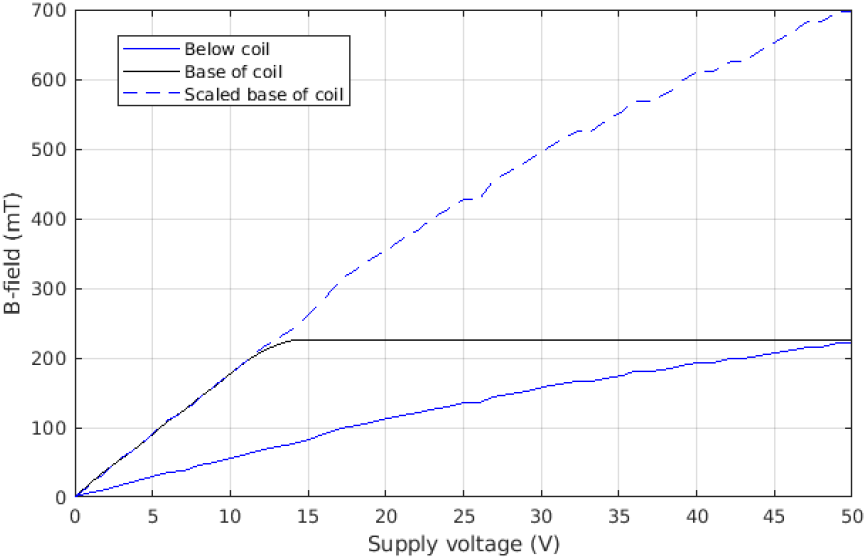
The maximum B-field against voltage. The black curve shows the B-field measured at the base of the coil; the Hall sensor saturates at about 14 V supply. The blue line shows the B-field measured about 3 mm below the coil. It reaches the saturation value for the sensor at about 50 V. The dashed blue line shows the B-field below the coil (blue line) scaled by a factor to match the low voltage measurements at the base of the coil. The continuation of this line at the higher voltages gives an indication of the B-field at the base of the coil at the higher voltages.

The time variation of the B-field is shown in Figure 8(a). The field reaches a peak after approximately 0.1 ms, and the pulse has decayed after 1 ms. The spatial distribution is shown in in Figure 8(b) and (c). The decay of the B-field with on-axis distance is shown in part (b). For these on-axis measurements, the B-field is expected to be purely axial and the Hall probe was oriented accordingly. At *z*=2 mm below the coil, the field intensity has decayed by around half compared with its value at the coil base. For a small magnetic dipole, we would expect a steep decay in field with increasing *z* of order 1/*z*^3^, but for the coil this would apply only at length scales greater than the size of the coil and thus this rapid decay would only be seen at greater distances than shown on the plot. Using the result of Figure 7, this suggests at 50 V supply, the B-field would be approximately 340 mT at 2 mm below the coil. The decay of the axial component *B_z_* of the *B*-field with radial displacement *r*, at an axial distance of *z*=2 mm below the coil, is shown in part (c). The field decays to about half its maximum 2 mm from the axis, showing significant focality. We note that at the larger radial distances the B-field will have a significant radial component, and thus the magnitude of the B-field will be larger than shown on this plot.

**Fig. 8.**
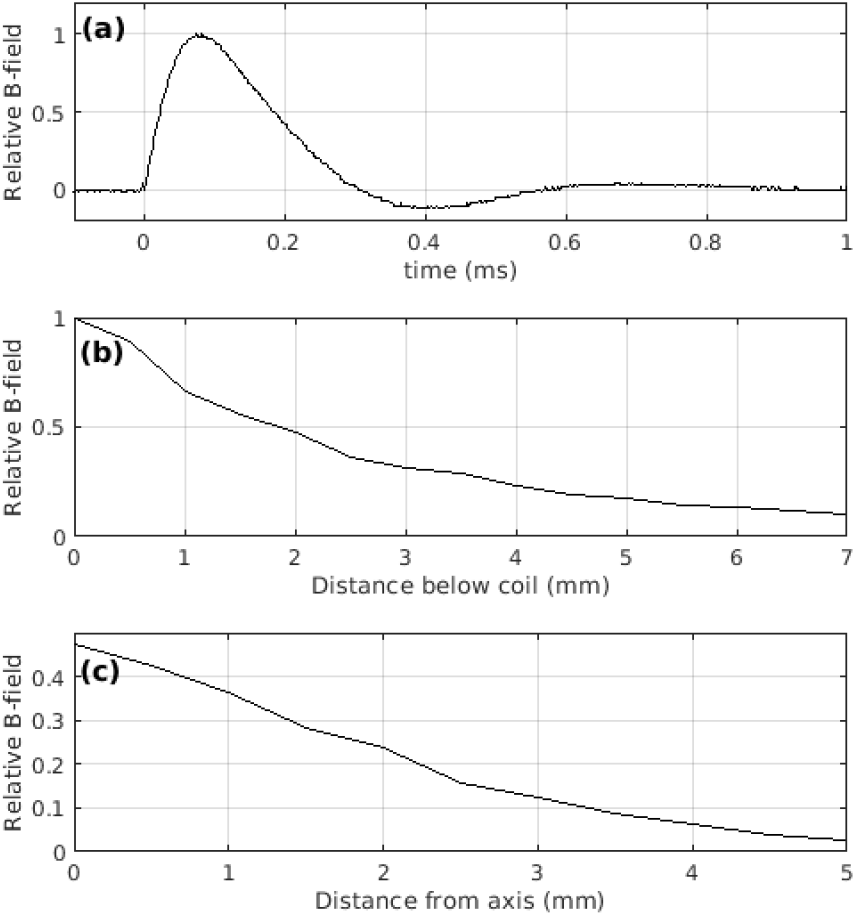
The temporal and spatial distribution of the B-field. (a) The time course of the B-field for a pulse measured at the base of the coil. (b) The B-field as a function of on-axis distance below the coil. (c) The axial component of the B-field as a function of radial distance from the axis of the coil measured at 2 mm below the base of the coil. All plots are taken at 12 V supply and are normalized to show a maximum relative B-field of unity at the base of the coil.

Figure 9 shows the induced electric field for 50 V supply. The *E*-field strength as a function of time is shown in Figure 9(a), for the position *z*=2 mm and *r*=6.5 mm. This position is selected since it gives the maximum E-field on a plane 2 mm below the coil. It shows a short pulse (around 0.1 ms duration) of *E*-field with a maximum strength of approximately 10 V/m occurring near instantaneously on the gating of the MOSFET, corresponding to the rapid increase in B-field of Fig. 8. A further negative peak occurs at 0.2 ms, when the decay in B-field is largest. The change of *E*-field with radial distance at *z*=2 mm below the base of the coil is shown in Figure 9(b). The plot shows a maximum *E*-field at 6.5 – 7 mm, and the E-field decays slowly with increasing radial distance. We emphasize that this measured *E*-field is due to the changing magnetic field generated by the coil, and any static electric fields are not able to be measured in this way. The measurement is in air; in brain tissue where charge separation may occur electric fields could be significantly different.

**Fig. 9.**
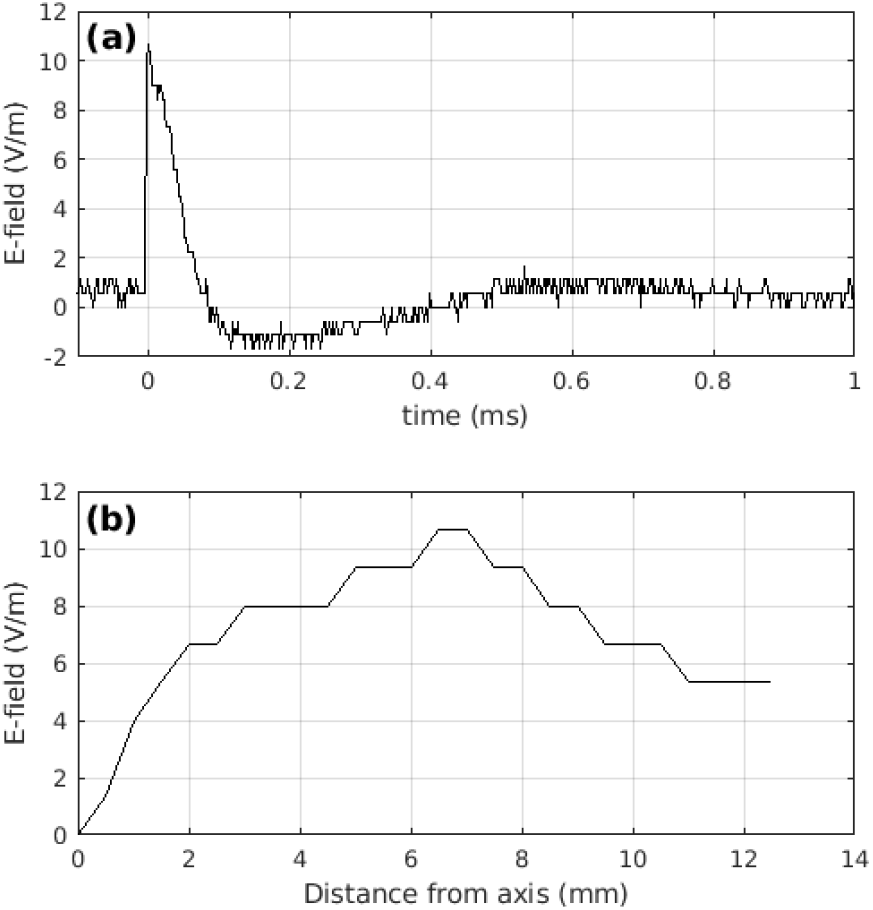
The E-field strengths as a function of time and space. (a) The Efield against time for a typical pulse at 50 V, 2 mm below the coil and at 6.5 mm radial displacement, measured with a wire loop and oscilloscope. (b) The peak E-field as a function of radial distance from the axis of the coil, measured at 2 mm below the coil at 50 V.

Figures 10 and 11 show the results of modelling with COMSOL Multiphysics for the *B*-and *E*-fields respectively, for 50 V supply. In Figure 10(a), the left white outline shows the powdered iron core geometry and the right white outline shows the coil geometry as described in Figure 2. Lines of B-field are shown, together with the magnitude of the B-field with a colour scale. Figure 10(b) shows a close-up in the vicinity of the base of the coil. The modelled *B*-field at the base of the coil, on axis, is around 1.7 T in magnitude, and 2 mm below the coil on axis it is about 600 mT. These values are around double the measured values of around 700 mT and 340 mT at the base of the coil and 2 mm below the coil respectively. The B-field is strongly focused at the corner of the taper (at *r*=1 mm, *z*=0 mm) meaning that the modelled B-field is stronger off axis than on axis. Figure 11 shows the Efield for the same situation. Part (a) shows the magnitude of the E-field on a colour scale. The mesh for the simulation is also shown. Part (b) shows the E-field in the vicinity of the base of the coil in more detail. The modeled maximum E-field at the base of the coil is around 15 V/m, with this maximum occurring about 1.5 mm from the axis. At 2 mm below the coil, the maximum E-field is around 8 V/m and occurs at about 4.5 mm from the axis. The modelled E-field is slightly smaller in magnitude and slightly more focused than measured experimentally. We have not investigated this discrepancy further but note that it might be a consequence of the unphysically sharp corners used in finite-element COMSOL approach, or uncertainties in the measurement process.

**Fig. 10.**
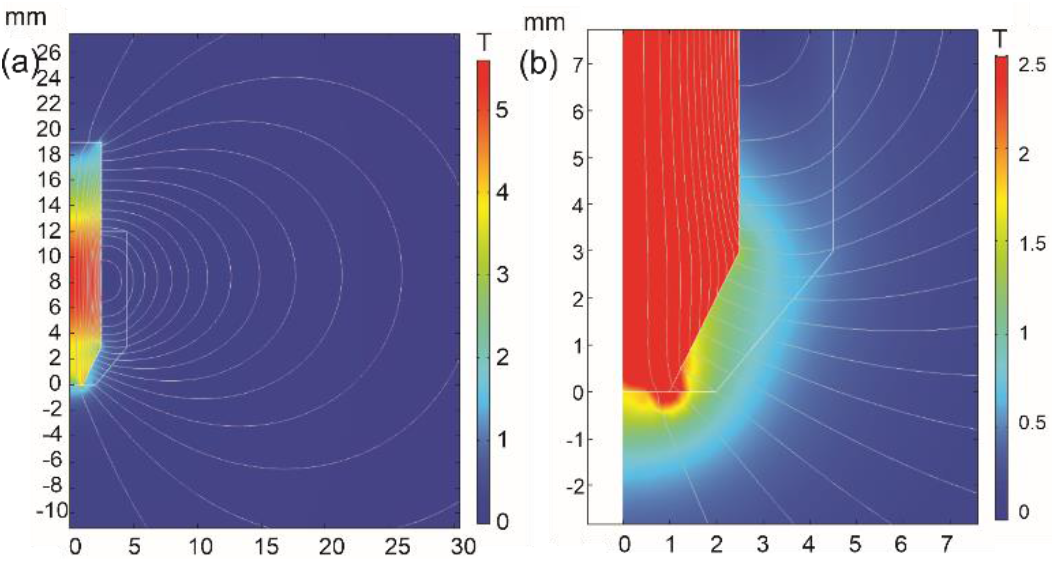
The B-field modelled with COMSOL Multiphysics using 2-D axial symmetry, at 50 V supply. (a) Lines of B-field, with magnitude indicated by the colour scale, in tesla. The geometry and dimensions (mm) of the core and coil are shown with the white outline. (b) A close-up of the region by the tip of the coil. Note the colour scale has been modified to better show the B-field immediately below the tip. In both parts (a) and (b) the horizontal axis is distance from the coil’s axis in mm and the vertical axis is the distance in mm along axis from the base of the coil.

**Fig. 11.**
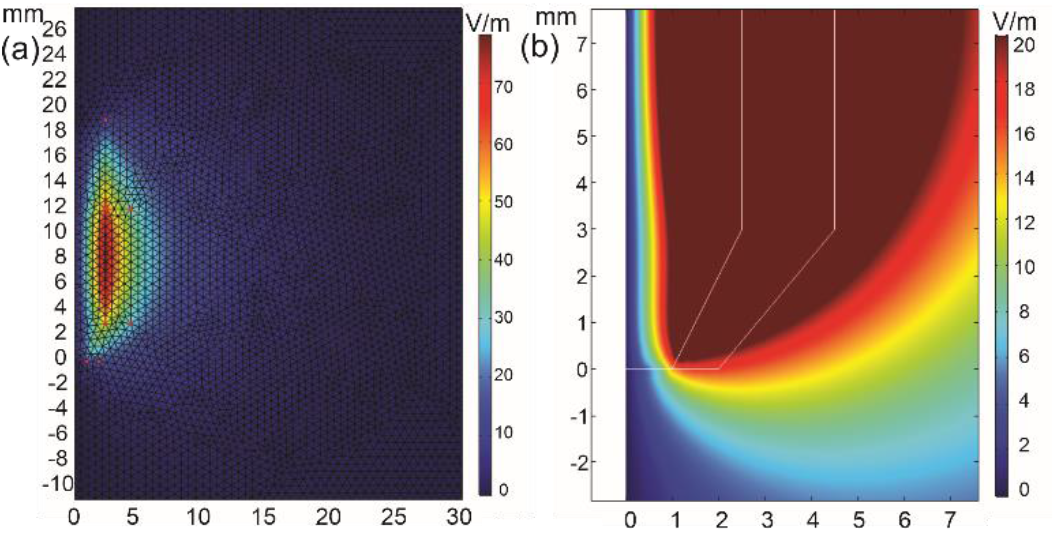
The E-field modelled with COMSOL Multiphysics using 2-D axial symmetry, for 50 V supply. (a) The magnitude of the E-field is shown by the colour scale, in V/m; its direction is azimuthal (i.e. perpendicular to the page). The figure also indicates the mesh used for the finite-element simulation. (b) A close-up of the region by the tip of the coil. The magnitude of the E-field is shown by the colour scale. The scale is different from part (a) in order to better show the variation in E-field in the vicinity of the tip of the coil. The core and coil geometry is marked by the white lines. The length scales are in mm for both parts (a) and (b).

### C. Temperature Rise

The heating of the coil was monitored for the case of the coil in air (i.e. not applied to the slice). For iTBS, the temperature of the coil rose by 10 degrees Celsius after 1200 pulses at 50 V power supply at room temperature of 22 degrees Celsius. For cTBS, the temperature of the coil rose by 20 degrees Celsius after 1200 at 50 V power supply at room temperature of 22 degrees Celsius. We have also recorded the temperature of the slice perfusion fluid during the experiments. It rose by just 0.2 Celsius after stimulation with 1200 pulses of cTBS (the same rise was also recorded after 1200 pulses of iTBS) with a background room temperature of 24.0 degrees Celsius. To test whether this increase in temperature affected SLE activity, we have independently measured SLE activity in a slice before and after raising the fluid temperature by 0.3 degrees Celsius through gentle heating. There was no change in SLE activity (results not shown) which suggests that any effect of temperature change in this experiment is likely to be insignificant.

### D. SLE Activity

We have measured the relative change in amplitude and frequency for each slice from its baseline. The relative change in amplitude from baseline after cTBS and sham stimulation is shown in Figure 12 for periods 0 - 10 min, 10 - 20 min and 20 - 30 min after stimulation. A similar plot for the relative changes in frequency is shown in Figure 13. Figures 14 and 15 show the results of iTBS stimulation on amplitude and frequency respectively. For each experiment, we applied a 2-way repeated measures ANOVA (RMANOVA) to test for changes due to treatment group (TBS or sham) and time. Where changes were statistically significant, we applied post-hoc t-tests to the distributions for the TBS and sham groups. The cut-off p-value for application of post-hoc t-test to the distributions are calculated as 0.0017 with the Bonferroni correction for 3 comparisons. For the changes in amplitude following cTBS, we obtained no statistical significance for either treatment group (*p* = 0.88) or time (*p* = 0.16). For changes in frequency following cTBS, we obtained a significant effect in both treatment group (*p* = 0.0053) and time (*p*= 0.024). For the changes in amplitude following iTBS, we see no effects either in treatment group (*p* = 0.060) or time group (*p* = 0.47); however, for changes in frequency we found a significant effect for the treatment group (*p* = 0.00094) but not for the time group (*p* = 0.098). Having established that the treatment group has an effect (the case for both cTBS and iTBS SLE frequency, but not SLE amplitude), we applied a post-hoc t-test to see where the effects were maximised. For frequency following cTBS, we see a significant decrease in SLEs inter-event frequency at 0-10 min (*p* = 0.00034) and 10-20 min (*p* = 0.0014) after stimulation, however, we see no change in 20-30 min after stimulation(*p* = 0.86). For frequency following iTBS, we see a significant increase in SLEs inter-event frequency at 0-10 min (*p* =0.00060) and 10-20 min (*p* = 0.0015) after stimulation, however, no change at 20-30 min after stimulation (*p*= 0.021).

**Fig. 12.**
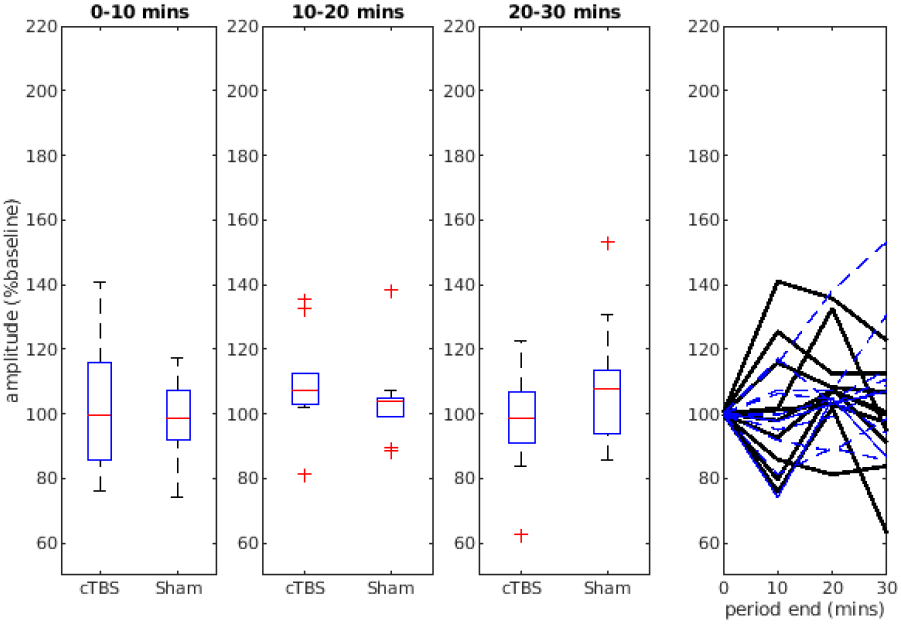
The relative change in SLE amplitude with respect to baseline for cTBS and sham stimulation for 0-10 min, 10-20 min and 20-30 min after stimulation. The fourth plot shows the response of the various individual slices; black denotes cTBS stimulation and blue-dashed sham.

**Fig. 13.**
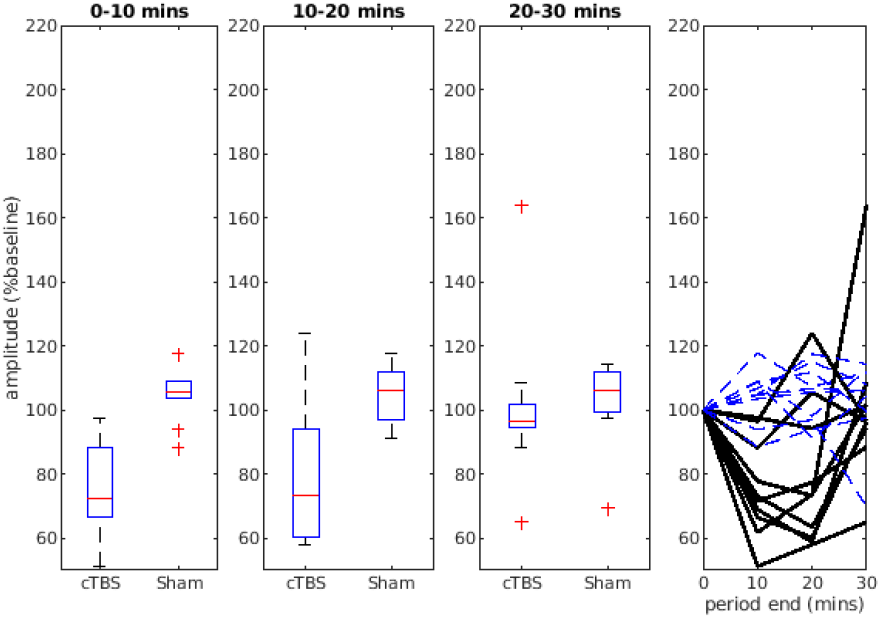
The relative change in SLE frequency after cTBS and sham stimulation for 0-10 min, 10-20 min and 20-30 min after stimulation. The fourth plot shows the response of the various individual slices; black denotes cTBS stimulation and blue-dashed sham.

**Fig. 14.**
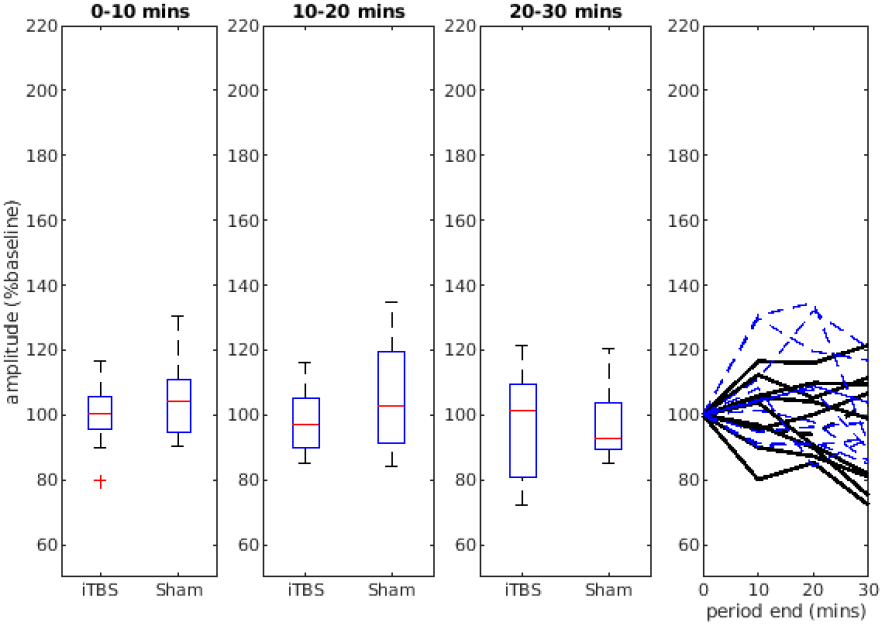
The relative change in SLE amplitude after iTBS and sham stimulation for 0-10 min, 10-20 min and 20-30 min after stimulation. The fourth plot shows the response of the various individual slices; black denotes iTBS stimulation and blue-dashed sham.

**Fig. 15.**
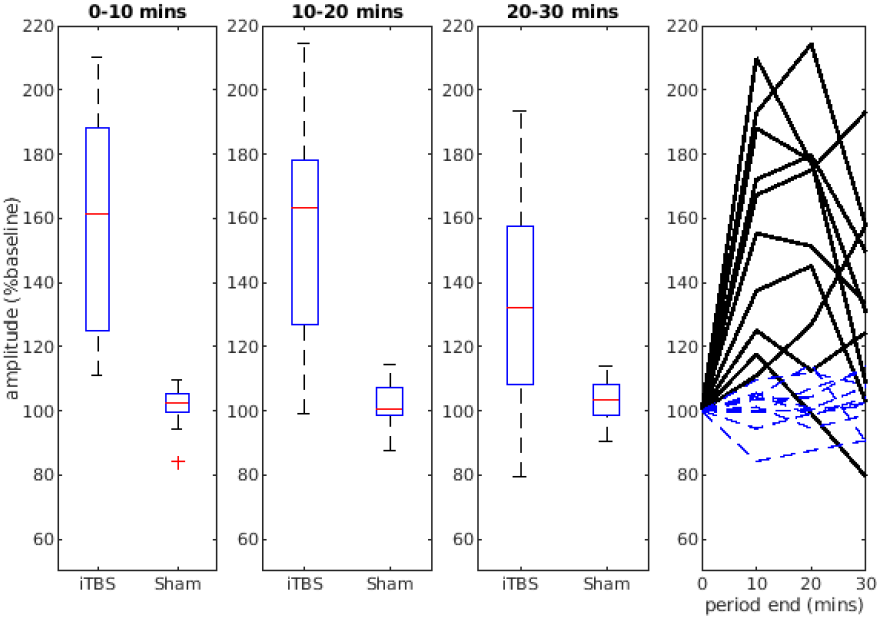
The relative change in SLE frequency after iTBS and sham stimulation for 0-10 min, 10-20 min and 20-30 min after stimulation. The fourth plot shows the response of the various individual slices; black denotes iTBS stimulation and blue-dashed sham.

## IV. Discussion

We have constructed a tapered mouse-specific TMS coil that exhibits stronger and more focused field strengths than many previously constructed small-scale coils [19, 28, 53, 63]. In particular, the B-field, of order hundreds of millitesla, compares favourably with previous coils. E-fields, of order 10 V/m, also compare favourably with previously constructed coils, but less notably than for B-field. Significant changes have been the reduction in the number of turns, the selection of a high saturation magnetization core material, and a tapering of one end of the core to focus the *B*-field and increase and focus the *E*-field. However, the field strengths, notably the *E*-field, are still much lower than the human TMS coils. Since we have measured and modeled fields in free-space, it is likely that the geometry of the brain surface could lead to localised increases in this value [28]. The region of strong E-field is annular in shape and its radius is significant when compared with the size of a mouse brain; experimentally the maximum E-field on a plane 2 mm below the base of the coil occurred at 6.5 mm from the axis. The modelled E-field was slightly more focused, with the maximum E-field when 2 mm below the coil occurring 4.5 mm from the axis. The coil design, having a significant thickness of turns around a central core, would need some modification before two such coils could be implemented together in a figure-of-eight style to further focus the field at a small point. There are several remaining challenges. Heating is a major problem in small-scale coils. This is because the reduction of a coil in radius and coil length will produce ten-fold increase in heating [19]. At 50 V power supply, the temperature rise of the coil after 1200 pulses of cTBS was 20 degrees celsius, with larger voltages expected to give significantly increased heating. For this coil to be suitable for *in vivo* experiments, there would need to be either cooling of the coil (e.g. through a heat sink) or thermal insulation to prevent conduction of heat from the coil to the head of the mouse. However, its application to experiments *in vitro* is still possible because of the removal of heat from the brain slice through forced convection by the aCSF. The primary source of heating is resistive heating in the coil. A permeable core ensures high enough inductance to give a sufficient ring-down time (larger than about 0.5 ms) in the inductor-capacitor circuit while keeping capacitance below a millifarad, thus reducing the pulse energy and heating [19]. However, this advantage is reduced when the core approaches saturation. It is possible that further improvement could be made by more sophisticated shaping of the core and coil. The winding of the coils could also be improved, to increase reproducibility. For example, the variation in resistance and inductance for three nominally similar coils was ±2% and ±20% respectively. Experimentally, these gave a variation in maximum B-field and maximum E-field of ±5% and ±20% respectively, which are similar in magnitude to uncertainties in the measurement.

We have demonstrated that the electromagnetic fields, although weaker than in the human case, are sufficient to cause biophysical changes in a brain slice over a time scale of 20 minutes when applied as cTBS or iTBS. The cTBS resulted in a decrease in frequency of SLE activity, whereas iTBS had an opposite effect. However, responses are highly variable between slices (Figures 13 and 15, final panels). These results are broadly consistent with the canonical decrease and increase in excitability following human cTBS and iTBS respectively [64, 65], although significant effects in our measurements were limited to 20 minutes after stimulation. While our method measures the local field potential due to spontaneous activity, the mechanisms are likely to be similar to those in humans, where long-term potentiation and long-term depression are reported to be largely NMDA-mediated phenomena [66]. No-magnesium SLE activity in cortical slices is NMDA-based [67] and lowering aCSF magnesium levels facilitates generation of LTP via an NMDA-dependent process in hippocampal [61] and neocortical [62] slices, So, while the experimental procedure has not been designed to test particular mechanisms of action of the coil on the brain tissue, it demonstrates that the fields are sufficient to achieve changes in the brain on the scale of several minutes. More discriminating experimental methods to assess plasticity and excitability of the cortex or other brain areas [68, 69] could be implemented, but these have challenges in terms of experimental set-up.

The coil construction is also challenging. Although the coil’s windings were secured with glue, the coil emitted audible clicks on discharge of the capacitor, especially at the higher voltages. Future designs will include better securing and insulating of the coil. Additionally, more sophisticated electronic engineering can be implemented to control the time-course of the current pulse and resistive heating.

## V. Conclusion

We have demonstrated a small TMS coil suitable for applying TMS to a mouse. The field strengths are larger than those reported by other authors using coils of similar focality. While the *B*-field strength at 50 V is a significant fraction of that produced by some human coils, the *E*-field strength is still substantially lower. However, we have demonstrated that such fields are still sufficient to change the biophysical behaviour of mouse cortex in-vitro with theta-burst stimulation, through an unspecified mechanism.

## Appendix

### A. Measurement of B-field at high intensities

In order to estimate the B-field at the base of the coil at 50 V supply (i.e. the maximum B-field in our experiments) using the same sensor, we compared the supply-voltage dependency of the B-field at the base of the coil with the supply-voltage dependency on axis 3 mm below the coil. That is, we assume *B*(*z, V*) = *f*(*z*) *g*(*V*), where *B* is the size of the B-field on axis (*r* = 0), *z* is the distance below the coil, *V* is the supply voltage and *f* and *g* are functions. We measured *B*(*z* = 0 mm, *V*) from 0 to 15 V supply at 1 V intervals, at which point the sensor reaches its maximum, and *B*(*z* = 3 mm, V) from 0 to 50 V supply. We then found the scale factor α such that *B*(*z* = 0 mm, *V*) = α *B*(*z* = 3 mm, *V*) over the range 0 to 15 V. We applied that same scale factor to the higher voltage measurements, thus estimating the B-field at the coil base at 50 V to be *α B*(z=3 mm, 50 V). Detailed results of this procedure are shown in Fig. 7. We emphasize that this measurement is indirect and the fields given at the higher voltages should be considered estimates. In particular, we note that any saturation of the core at lower voltages would lead to an overestimate of field strength at the higher voltages. However, the powder iron core has a higher saturation magnetization (1.5 T) than our field strengths, so this is unlikely to have a large effect.

### B. Measurement of electric field

We now describe the electric field measurements in more detail. Figure 16 shows the geometry of the coil of radius and the wire loop used for measurement of the *E*-field. In the case shown, the loop is being used to measure the field at a distance *h* below the coil, at a distance *r* from the axis. The loop is oriented so that its apex lies on the axis of the coil, and its short side ‘b’ lies parallel to the direction of the induced electric field, which is azimuthal for a cylindrically symmetric situation. The long sides ‘a’ and ‘c’ are thus perpendicular to the induced electric field. The voltage induced around the loop is equal to the line integral of the electric field around the loop. The two long sides contribute zero to this integral, being perpendicular to the induced electric field, whereas the short side contributes *E b*, where *E* is the *E*-field and *b* is the length of the side. Thus by measuring the induced voltage *V* we find the induced electric field as *E* = *V/b*. Such an arrangement is only applicable to measuring the induced electric field in a cylindrically symmetric geometry. To measure in different geometries (e.g. of a figure-of-eight coil) or to measure static electric fields, more complicated measurement techniques are required.

**Fig. 16.**
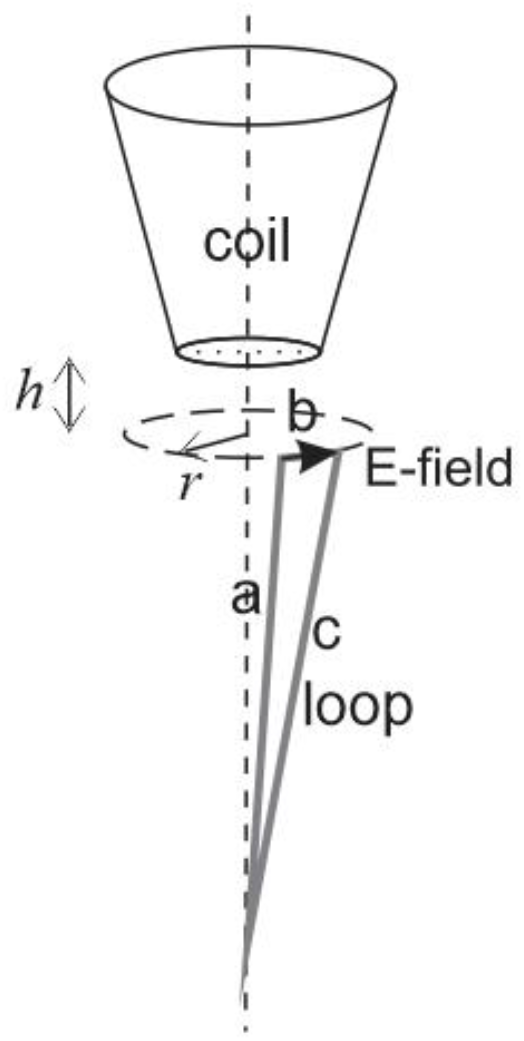
The orientation of the coil and core for measurement of induced electric field at a distance h below the coil and distance r from the axis of the coil. The apex of the loop is located on the coil axis, so that the wires ‘a’ and ‘c’ are perpendicular to the electric field. Only the short wire ‘b’ contributes to the voltage induced around the loop.

### C. Preparation of brain tissue and electrophysiological measurement

We prepared 10 coronal brain slices from 5 mice (2 slices from each mouse) for iTBS experiments and 10 coronal brain slices from 3 mice (3 or 4 slices from each mouse) for cTBS experiments. Mice were of both sexes with the genetic background C57. Their age varied from 4 to 10 months old. They were housed in a temperature-controlled room with unlimited access to food and water. The mice were anesthetized with carbon dioxide and decapitated. The brain was dissected and placed in ice-cold HEPES-buffered ‘normal’ artificial cerebrospinal fluid (aCSF), oxygenated with 95% oxygen (Perfecto2 oxygen concentrator, Invacare, New Zealand). HEPES (4-(2-hydroxyethyl)-1-piperazineethanesulfonic acid) is a zwitterionic sulfonic acid buffering agent. The farmost posterior and anterior coronal sections of the brain were removed with a razor blade. The remaining brain (approximately between Bregma 1 and −5 mm) was glued onto a metallic plate, placed into oxygenated ice-cold HEPES-buffered ‘normal’ aCSF, and coronally sectioned into slices 400 microns thick (Vibratome, Campden Instruments Ltd., United Kingdom). HEPES ‘normal’ artificial cerebrospinal fluid contained 130 mM sodium chloride, 2.5 mM potassium chloride, 1 mM magnesium chloride, 2 mM calcium chloride, 2.5 mM NaHCO_3_, 10 mM HEPES and 20 mM D-glucose in double distilled water. The aCSF was oxygenated with 95% oxygen (Perfecto2 oxygen concentrator, Invacare, New Zealand). After the brain slices were sectioned, they were moved into oxygenated HEPES-buffered ‘no-magnesium’ aCSF for a minimum 1-hour recovery at room temperature. The ‘no magnesium’ aCSF contained 130 mM sodium chloride, 5 mM potassium chloride, 2 mM calcium chloride, 2.5 mM NaHCO_3_, 10 mM HEPES and 20 mM D-glucose. The pH level of all solutions was adjusted to 7.4 with 10 M sodium hydroxide. Apart from HEPES (ITW Reagents, Spain) and sodium chloride (EMSURE, Denmark), the aCSF ingredients were obtained from Sigma (USA).

Following the minimum recovery period, one slice at a time was moved to a submersion-style perfusion bath (Kerr Scientific Instruments, New Zealand). The perfusion bath was replenished continuously with oxygenated ‘no-magnesium’ aCSF by gravity-feed at a rate of 5 ml/min. We clamped the coil above the perfusion bath where the slice was resting. We positioned a 75 micron diameter silver/silver chloride electrode (GoodFellow Ltd., United Kingdom) in layer IV of the mouse cortex (Figure 4) to record spontaneous local field potential (LFP) activity. The analog signal was recorded via a headstage placed in close proximity to the slice preparation. The signal was amplified 1000 times, low pass (300 Hz) and high pass (1 Hz) filtered (Model 3000 differential amplifier, A-M Systems, USA) and converted to a digital signal (Power-lab, ADInstruments, Australia). The amplified and filtered signal was stored for later analysis. The temperature of the aCSF was monitored with a thermocouple probe.

